# Identification of Viral and Cellular Proteins in Proximity to the HSV-2 pUL16 Tegument Protein in Infected Cells

**DOI:** 10.64898/2026.07.07.737067

**Authors:** Safara M. Holder, Alexis Lubinsky, Maike Bossert, Bruce W. Banfield

## Abstract

Orthologs of the herpes simplex virus (HSV) pUL16 tegument protein are conserved throughout the *Orthoherpesviridae* family. During HSV infection, pUL16 functions in the nuclear egress of nascent nucleocapsids from the nucleus to the cytoplasm, prevents the docking of nascent cytoplasmic nucleocapsids to nuclear pore complexes, promotes the final envelopment of cytoplasmic nucleocapsids, and enhances cell-to-cell spread of virus infection. How pUL16 performs these diverse functions is poorly understood. To gain further insight into the mechanisms by which pUL16 mediates its activities, we utilized a BioID approach to identify cellular and viral proteins in proximity to pUL16 during the infection of human keratinocytes. By comparing proteins in proximity to pUL16 during infection with proteins in proximity to its well-known virus-encoded binding partner, pUL21, we provide new insight into the activities of pUL16 that likely occur in complex with pUL21 and those that are independent of pUL21. A key function of pUL21 is to deliver protein phosphatase 1 (PP1) to viral and cellular substrates to mediate their dephosphorylation. Intriguingly, the findings presented suggest that pUL16 interactions with pUL21 may regulate the isoform of PP1 that is bound to pUL21 and thereby regulate the specificity of substrate dephosphorylation.

**Importance:** HSV-1 and HSV-2 are important human pathogens that currently infect roughly 3.8 billion and 520 million people, respectively. These viruses cause lifelong, recurrent, infections and cause a variety of diseases including vesicular lesions of the oral and genital mucosa, corneal blindness, meningitis, encephalitis and devastating neonatal infections. The HSV pUL16 tegument protein performs a number of critical functions for the virus that influence virion assembly and the spread of infection between cells. In this study we have identified cellular and viral proteins that are proximal to pUL16 during infection of human keratinocytes, providing new insight into the mechanisms used by pUL16 to perform its activities.

## Introduction

Herpes simplex viruses (HSVs) are important and ubiquitous human pathogens that establish lifelong infections and cause a wide range of human diseases, ranging from trivial asymptomatic infections to life-threatening disseminated neonatal disease. HSV virions are complex assemblages, with infectious particles consisting of an icosahedral nucleocapsid, containing a linear double-stranded DNA genome, surrounded by a lipid envelope embedded with glycoproteins (1). Between the nucleocapsid and the envelope lies a proteinaceous compartment called the tegument, which encloses roughly 20 virus-encoded proteins and approximately 50 host cell-encoded proteins (2).

The initial stages of herpesvirus assembly take place in the nucleus, where newly replicated virus genomes are packaged into preformed capsids. DNA-containing nucleocapsids gain access to the cytoplasm by undergoing a vesicular transport process through the nuclear envelope called nuclear egress. Once in the cytoplasm, the tegument is formed through the recruitment of tegument proteins to capsid components, interactions among tegument proteins, and interactions between tegument proteins and the cytoplasmic tails of membrane glycoproteins destined for the envelope of mature virions (3). The virion acquires its final envelope through budding of capsid-tegument complexes into membranes derived from a post-Golgi compartment, a process referred to as secondary envelopment. Vesicles containing enveloped virus then traffic to, and fuse with, the plasma membrane of the cell, releasing virus into the extracellular environment, which can go on to infect new cells, or new hosts (4, 5). In addition to infection by cell-free extracellular virions, cells can become infected through contact with adjacent infected cells by a distinct process called cell-to-cell spread.

Tegument proteins perform many different functions during HSV infection. Immediately after fusion of the virion envelope with a host cell membrane, capsid-associated tegument proteins facilitate the recruitment and regulation of microtubule motors that mediate the transport of the capsid from the cell periphery towards the nuclear envelope, where genome uncoating takes place (6–13). Many other tegument proteins dissociate from the capsid after its delivery to the cytoplasm and provide important functions, including the shutoff of host cell protein synthesis, suppression of host innate immune responses, and initiation of viral immediate early gene expression (8, 14–17). Later in infection, tegument proteins function in multiple aspects of virion assembly and modulate host processes that facilitate virus replication, while inhibiting those that restrict it.

Our laboratory, and several others, have been studying the function of the HSV tegument protein pUL16 (18). In HSV-1 and HSV-2, pUL16 functions in the nuclear egress of nucleocapsids, the secondary envelopment of nucleocapsids at cytoplasmic membranes, prevention of nascent cytoplasmic nucleocapsid binding to nuclear pore complexes, and the cell-to-cell spread of virus infection (19–27). This latter process is mediated through interactions of pUL16 with the membrane-associated tegument protein pUL11, the capsid-associated tegument protein pUL21, and the viral membrane glycoprotein, gE (28–31). Additionally, HSV-1 pUL16 localizes to mitochondria and appears to promote mitochondrial activity through its interaction with the ADP/ATP translocase adenine nucleotide transporter 2 (ANT2) (32, 33).

To gain a better understanding of the many functions of HSV-2 pUL16 during viral replication, we performed proximity-dependent biotinylation (BioID) experiments in human keratinocytes (HaCaT) to identify cellular and viral proteins in proximity to pUL16 during HSV-2 infection. To do this, we constructed a recombinant HSV-2 strain in which pUL16 was fused to the promiscuous biotin ligase miniTurbo (mT; 16-mT) (34). When infected cells expressing the pUL16mT-fusion are incubated with biotin, mT produces a short-lived reactive biotin species that becomes covalently linked to proximal lysine residues within an approximately 10 nm radius of the mT moiety (35). Cells are lysed and biotinylated proteins are captured on streptavidin-conjugated beads that, due to the strength of the biotin-avidin interaction, can withstand washing with buffers containing salts and detergents at concentrations that interfere with both specific and non-specific protein-protein interactions. Thus, the BioID approach has the advantage of a low non-specific background, improving confidence in the identified proteins. Additionally, BioID can detect weak or transient interactions between bait and prey proteins that would not be identified by co-affinity purification methodologies. Importantly, BioID offers the advantage of probing proximal protein interactions in the context of living cells. Our findings indicate that 21 cellular proteins and 4 viral proteins were in proximity to pUL16mT between 8 and 10 hours post-infection (hpi).

## Results and Discussion

### Analysis of an HSV-2 strain expressing pUL16 fused to the biotin ligase miniTurbo (mT)

To enable the identification of proteins in proximity to pUL16 during infection, we constructed an HSV-2 186 strain that expresses pUL16 fused to the promiscuous biotin ligase, mT (34, 36), at the UL16 locus using *en passant* mutagenesis of an HSV-2 186 BAC (37, 38). This strain, 16-mT, replicated to a somewhat lower level (p=0.0647) in human keratinocytes (HaCaT) than the wild-type (WT) 186 parental strain and at a significantly higher level (p=0.0041) than the UL16 null strain, Δ16 (21) (Figure 1A). 16-mT formed plaques on HaCaT cells that were intermediate in size between the large WT plaques and the small plaques formed by Δ16 in both the presence and absence of exogenous biotin (Figure 1B, C). Taken together, these results indicate that the pUL16mT fusion protein maintains activities that support virus replication and spread, and that 16-mT is suitable for the identification of pUL16 proximal proteins.

**Figure 1.**
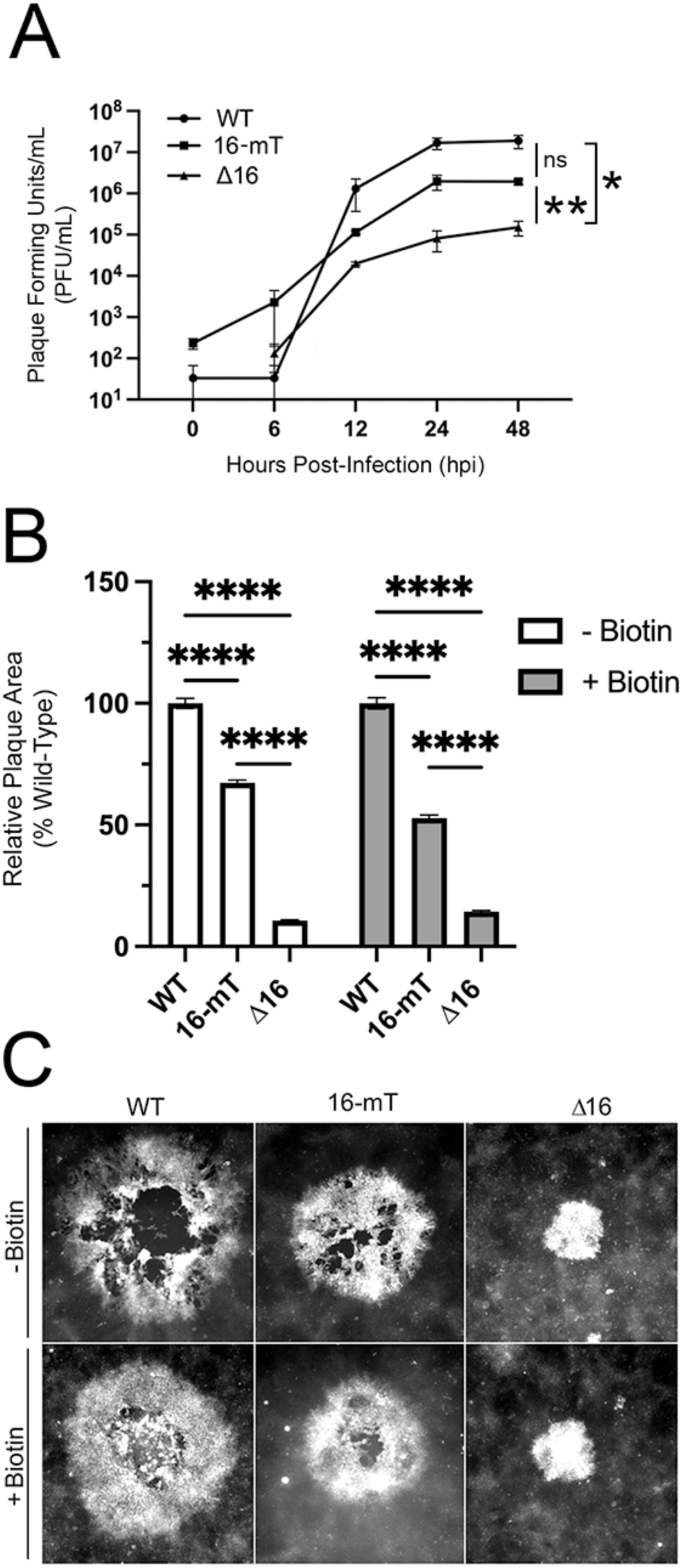
Analysis of 16-mT. **(A)** Replication analysis. HaCaT cells were infected with HSV-2 186 (WT), HSV-2 pUL16mT (16-mT), and HSV-2 ΔUL16 (Δ16) in triplicate at an MOI of 1. Following a one-hour inoculation period, the inoculum was removed and extracellular virus was inactivated by briefly incubating the cells in a low-pH citrate buffer prior to the addition of fresh medium. At 6, 12, 24, and 48 hours post-infection, cells were scraped into the medium and subjected to three rounds of freezing (-80°C) and thawing (37°C) to release virus from infected cells. Infectious virus was quantified by plaque assay on monolayers of Vero16 cells. Error bars represent the standard error of the mean from biological triplicates (*p<0.05. **p<0.01). **(B)** Plaque size analysis of WT, 16-mT, and Δ16 in the absence and presence of exogenous biotin (50 µM). At 48 hours post-infection, HaCaT cells were fixed, permeabilized, and stained for the HSV-2 protein, pUs3. The average plaque area of each strain relative to WT is shown (n=55 for each strain). Error bars represent the standard error of the mean (****, p<0.0001). **(C)** Representative images of plaques produced on HaCaT cells by each virus strain in the presence and absence of 50 µM exogenous biotin.

Vero cells were either mock infected, infected with an HSV-2 strain that expresses pUL21 fused to mT, 21-mT (39), or 16-mT and were treated with 50µM biotin for two hours at 18 hours post-infection (hpi) prior to the preparation of whole cell lysates. We included the well-characterized 21-mT strain in this analysis as a positive control for protein biotinylation (39). Biotinylated proteins from cell lysates were visualized on western blots using streptavidin-conjugated horse radish peroxidase (SA-HRP) and a chemiluminescent substrate (Figure 2A). Additionally, expression of pUL16 and mT fusion proteins was detected using antisera against pUL16 and BirA*, respectively. As expected, predominantly endogenous biotinylated mitochondrial proteins of roughly 130 kDa (pyruvate carboxylase) and 80 kDa (propionyl-CoA carboxylase alpha chain and methylcrotonoyl-CoA carboxylase subunit alpha) were identified in mock-infected lysates and in lysates from infected cells that were not treated with biotin (40). Many biotinylated infected cell proteins were observed in lysates prepared from 21-mT infected cells that were treated with biotin. By comparison, fewer biotinylated proteins were detected in cells infected with 16-mT after biotin treatment than in 21-mT infected cells, but more than what was observed in mock infected cells. Whereas WT pUL16 and the pUL21mT fusion protein were detected in extracts prepared from 21-mT infected cells, WT pUL16 was not detected in 16-mT infected cells, as expected; instead, these cells expressed pUL16mT, which was detected with both pUL16 and BirA* antisera.

**Figure 2.**
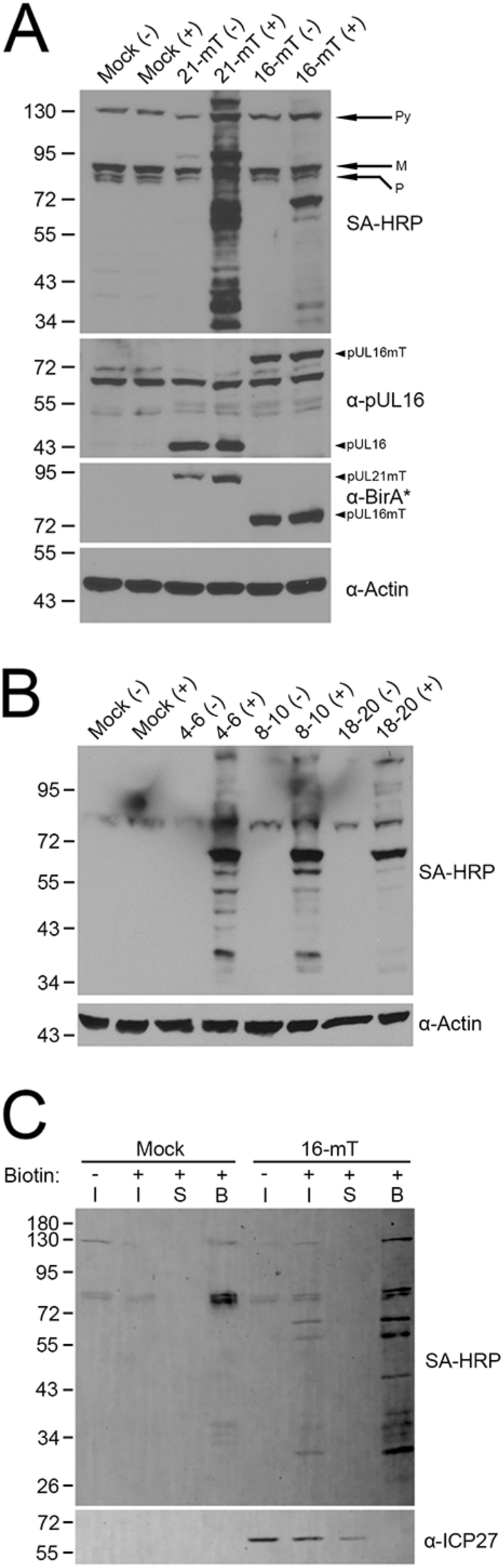
Biotinylation profile of HSV-2 pUL16mT. **(A)** pUL16mT is capable of biotinylating infected cell proteins. Vero cells were mock-infected, infected with HSV-2 pUL21mT (21-mT), or HSV-2 pUL16mT(16-mT) at an MOI of 1, and treated with 50 µM exogenous biotin at 18 hpi for a period of two hours (+). No exogenous biotin was added to the control samples (-). Whole cell lysates were prepared and analyzed by western blot using streptavidin-HRP (SA-HRP) to visualize biotinylated proteins and antibodies against pUL16, BirA* (reactive against mT), and actin. Arrows on the right side of the SA-HRP blot indicate the migration positions of the endogenously biotinylated proteins pyruvate carboxylase (Py), methylcrotonoyl-CoA carboxylase subunit alpha (M) and propionyl-CoA carboxylase alpha chain (P). Arrowheads on the right side of the ⍺-pUL16 and the ⍺-BirA* blots indicate the migration positions of pUL16mT, pUL16 and pUL21mT. **(B)** Time course of pUL16mT biotinylation. HaCaT cells were mock-infected or infected with 16-mT at an MOI of 1 and treated with 50 µM exogenous biotin (+) at 4, 8, or 18 hpi for a period of 2 hours (4-6, 8-10, or 18-20, respectively). No exogenous biotin was added to the control samples (-). Whole cell lysates were prepared and analyzed by western blotting using SA-HRP to visualize biotinylated proteins and antibodies against actin as a loading control. **(C)** Affinity purification of proteins biotinylated by pUL16mT between 8-10 hpi. HaCaT cells were mock-infected or infected with 16-mT at an MOI of 1 for 8 hours and treated with 50 µM exogenous biotin for 2 hours. Biotinylated proteins were affinity-purified using streptavidin-Sepharose beads. Following incubation, bead-bound proteins were pelleted by centrifugation (Bound; B), and post-pull-down supernatants (S) were collected. A portion of each sample was retained prior to the addition of streptavidin-Sepharose beads (Input; I). Biotinylated proteins were visualized by western blotting using SA-HRP. As an infection control, samples were also probed for the HSV-2 protein ICP27. Migration positions of molecular weight markers (kDa) are shown on the left side of each blot.

To determine the optimal time after 16-mT infection for maximal biotinylation of pUL16mT proximal proteins, a time-course experiment was performed. HaCaT cells were mock-infected or infected with 16-mT for 4, 8 and 18 hours and then treated (+), or not treated (-), with 50 µM biotin for two hours prior to the preparation of cell lysates. Biotinylated proteins in cell lysates were visualized by western blotting using SA-HRP (Figure 2B). Many more biotinylated protein species were detected when biotin labelling occurred between 4-6 hpi and 8-10 hpi than from 18-20 hpi. Based on these findings, coupled with pUL16mT being expressed as a late protein, the 8-10 hpi biotin labelling protocol was used in subsequent experiments.

Biotinylated proteins were efficiently purified using streptavidin-conjugated Sepharose beads, as evidenced by the lack of biotinylated proteins in the post-pull-down supernatants (Figure 2C). The subcellular localization of proteins biotinylated by pUL16mT between 8 and 10 hpi was determined by staining HaCaT cells with Alexa Fluor 488-labelled streptavidin (SA-488) (Figure 3). Whereas weak SA-488 signals were observed in mock and infected cells in the absence of biotin and in mock-infected cells treated with 50 µM biotin for two hours, robust pancellular labelling of biotinylated proteins was observed in biotin-treated 16-mT infected cells. Consistent with the reported localization of pUL16 to the nuclear rim (27), biotin labelling of the nuclear envelope was readily apparent in 16-mT infected cells (arrowheads) as evidenced by co-localization with the viral pUL34 protein that localizes predominantly to the inner nuclear membrane (INM) (41).

**Figure 3.**
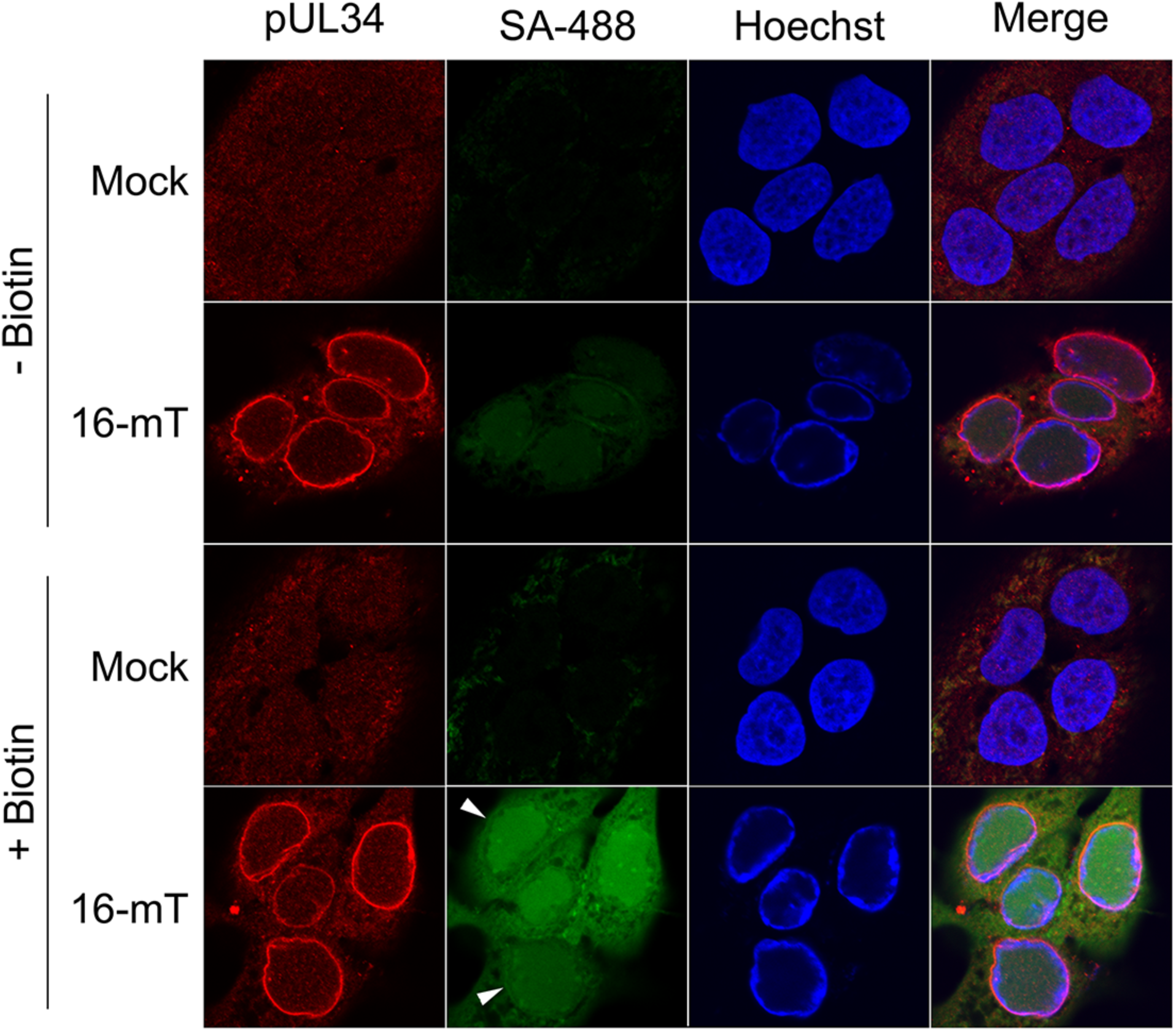
Subcellular localization of proteins biotinylated by pUL16mT at 10 hours post-infection. HaCaT cells were mock-infected or infected with 16-mT at an MOI of 1. At 8 hpi, cells were treated with 50 µM exogenous biotin for two hours, fixed, permeabilized, and stained with antisera against the viral protein pUL34 (red), streptavidin-conjugated Alexa Fluor 488 (SA-488; green), and the DNA stain Hoechst 33342 (blue). Arrowheads point to SA-488 labelling of the nuclear envelope. The scale bar is 10 µm and applies to all panels.

### Identification of proteins biotinylated by 16-mT from 8-10 hpi

HaCaT cells were infected at an MOI of 3 in triplicate with 16-mT for 8 hours, treated with or without 50 µM biotin for two hours, after which biotinylated proteins were affinity-purified on streptavidin-conjugated Sepharose beads. Proteins on beads were digested with trypsin and resulting peptides were identified by LC-MS/MS. As a specificity control, HaCaT cells were also infected in triplicate with EGFP-mT, an HSV-2 recombinant strain that expresses EGFP fused to mT from the Us2 locus for 8 hours and incubated with or without 50 µM biotin for two hours. Similarly, biotinylated proteins from EGFP-mT infected cells were affinity-purified using streptavidin-Sepharose beads and identified by LC/MS/MS. Raw mass spectrometry data are shown in the Supplementary Data 1 file.

After removing common contaminants (e.g. keratins) and contaminants frequently associated with BioID experiments (e.g. filamin A) (42), triplicate experiments were normalized based on spectral counts obtained from endogenously biotinylated proteins pyruvate carboxylase, propionyl-CoA carboxylase alpha chain and methylcrotonoyl-CoA carboxylase subunit alpha. Spectral counts from triplicate samples were averaged and values obtained from the “minus biotin” samples were subtracted from their corresponding “plus biotin” samples. Biotinylated proteins were then ranked based on the percent coverage of the peptides identified for a given protein, and proteins with less than 5% coverage were excluded. Finally, the normalized, averaged spectral counts of pUL16mT biotinylated proteins were divided by the normalized, averaged spectral counts of the same EGFP-mT biotinylated protein to yield a “fold enrichment over EGFP-mT” value. pUL16mT proteins with less than 2-fold enrichment over EGFP-mT were excluded. Based on these criteria, 21 cellular proteins and 4 viral proteins were further analyzed and are listed in Table 1.

**Table 1.** Viral and Cellular Proteins Proximal to pUL16 from 8-10 hpi Ranked by Normalized Percent Coverage.

| Protein Name | Gene Name | Molecular Weight | <sup>1</sup>Normalized Spectral Counts | <sup>2</sup>Fold-Enrichment Over EGFP-mT | <sup>3</sup>Normalized Percent Coverage <sup>4</sup>(±SD) |
| --- | --- | --- | --- | --- | --- |
| HSV-2 Tegument Protein pUL21 | TEG4_HHV2H | 57 kDa | 40.1 | 20.8 | 43.7 (±1.2) |
| Lamina-associated polypeptide 2, isoforms beta/gamma | LAP2B | 51 kDa | 22.9 | 2.7 | 42.0 (±4.4) |
| Transcription factor jun-B | JUNB | 36 kDa | 14.7 | 2.1 | 31.7 (±10.5) |
| Virion host shutoff protein (pUL41) | SHUT_HHV2H | 55 kDa | 11.8 | 3.6 | 27.6 (±5.5) |
| Cytoplasmic envelopment protein 2 (pUL16) | CEP2_HHV2H | 40 kDa | 9.0 | 20.1 | 25.0 (±3.6) |
| Gamma-interferon-inducible protein 16 | IFI16 | 88 kDa | 29.2 | 2.4 | 22.7 (±7.8) |
| Arf-GAP domain and FG repeat-containing protein 1 | AGFG1 | 58 kDa | 12.1 | 4.3 | 18.3 (±2.9) |
| Synaptotagmin-like protein 4 | SYTL4 | 76 kDa | 13.8 | 2.5 | 17.3 (±11.0) |
| Protein FAM83H | FAM83H | 127 kDa | 19.0 | 2.4 | 16.4 (±7.2) |
| Serine/threonine-protein phosphatase PP1-gamma catalytic subunit | PPP1CC | 37 kDa | 6.2 | 41.0 | 15.8 (±8.5) |
| Lamina-associated polypeptide 2, isoform alpha | LAP2A | 75 kDa | 16.0 | 2.2 | 15.7 (±1.2) |
| 28S ribosomal protein S31, mitochondrial | MRPS31 | 45 kDa | 6.3 | ∞ | 15.0 (±4.0) |
| Emerin | EMD | 29 kDa | 4.7 | 6.4 | 12.7 (±0.6) |
| Zinc finger CCCH-type antiviral protein 1 | ZCCHV | 101 kDa | 9.1 | 2.2 | 12.6 (±8.2) |
| Inner nuclear membrane protein Man1 | MAN1 | 100 kDa | 9.5 | 3.0 | 12 (±5.5) |
| Tubulin beta chain | TUBB | 50 kDa | 3.9 | ∞ | 11.7 (±1.2) |
| Delta(14)-sterol reductase LBR | LBR | 71 kDa | 11.3 | 4.2 | 10.8 (±1.4) |
| ATP-dependent RNA helicase DDX3X | DDX3X | 73 kDa | 6.6 | 2.3 | 8.9 (±3.6) |
| AP-4 complex subunit epsilon | AP4E1 | 127 kDa | 9.4 | 12.7 | 8.8 (±3.7) |
| Heterogeneous nuclear ribonucleoproteins C1/C2 | HNRNPC | 34 kDa | 3.9 | 5.2 | 8.6 (±5.4) |
| Golgin subfamily A member 4 | GOLGA4 | 261 kDa | 17.2 | 6.9 | 8.3 (±7.9) |
| Inner tegument protein pUL37 | ITP_HHV2H | 119 kDa | 9.0 | 3.6 | 8.0 (±5.6) |
| Protein ELYS | ELYS | 253 kDa | 14.7 | 2.1 | 7.2 (±4.0) |
| Protein CLEC16A | CLEC16A | 118 kDa | 6.0 | ∞ | 5.6 (±4.3) |
| Phosphatidylinositol-binding clathrin assembly protein | PICALM | 71 kDa | 4.7 | 8.1 | 5.5 (±2.5) |

<sup>1</sup>Spectral counts from three biological replicates were normalized to endogenously biotinylated cellular proteins then averaged. Averages of the no-biotin control samples were subtracted from the plus-biotin experimental samples to determine the normalized spectral count.
<sup>2</sup>Normalized spectral counts of averaged plus-biotin 16-mT spectral counts divided by the averaged plus-biotin EGFP-mT spectral counts.
<sup>3</sup>Average of the percent coverage of three biological replicates of the no-biotin control samples was subtracted from the average of the percent coverage of the plus-biotin experimental samples to determine the normalized percent coverage.
<sup>4</sup>Standard deviation (SD) of the three plus-biotin biological replicates

### Analysis of proteins in proximity to both pUL16 and pUL21

The top-ranked protein in proximity to pUL16 was the HSV-2 tegument protein, pUL21, a well-characterized interaction partner of pUL16 that served to validate our approach (30). Additionally, the cellular serine/threonine protein phosphatase PP1 gamma, which binds directly to pUL21 was a top hit, suggesting that pUL16, pUL21, and PP1 gamma are contained within the same complex in HSV-2 infected cells. pUL21 is a PP1 adapter protein that targets PP1 phosphatase activity to both viral and cellular substrates (43, 44). How pUL16 influences/directs the activity of PP1 gamma in pUL16/pUL21/PP1 gamma complexes is unclear at present. Interestingly, our recent report that identified proteins in complex with pUL21 in infected cells detected similar levels of two PP1 isoforms, PP1 alpha and PP1 gamma, in complex with, and in proximity to, HSV-2 pUL21 (39). Here, only the PP1 gamma isoform was identified in proximity to pUL16, raising the intriguing possibility that pUL16 is found specifically in PP1 gamma-containing complexes. In support of this idea, it is notable that our recent pUL21 protein proximity analysis failed to identify pUL16 as a top hit, suggesting that the majority of pUL21 in the infected cell is not in complex with pUL16 (39).

Fifteen of the 25 proteins in proximity to pUL16 were previously shown to be in proximity to pUL21 (Figure 4) (39). Of these, emerin, LAP2 beta/gamma, lamin B receptor (LBR), and MAN1 are well-characterized INM proteins that have been proposed to recruit pUL21 to the INM (39). As pUL21 is required to localize pUL16 to the nuclear envelope (27), these findings suggest that pUL16 is proximal to these molecules by virtue of its interaction with pUL21. Both pUL21 and pUL16 are important for the nuclear egress of nascent HSV nucleocapsids from the nucleoplasm to the cytoplasm and thus their recruitment to the INM may facilitate their role in this process (20–23, 37). Similarly, the RNA helicase DDX3X, which was also found in proximity to both pUL16 and pUL21 (Table 1, Figure 4, (39)), functions in nuclear egress and is concentrated at the INM during HSV-1 infection, likely through its interaction with the HSV nuclear egress complex component pUL31 (45).

**Figure 4.**
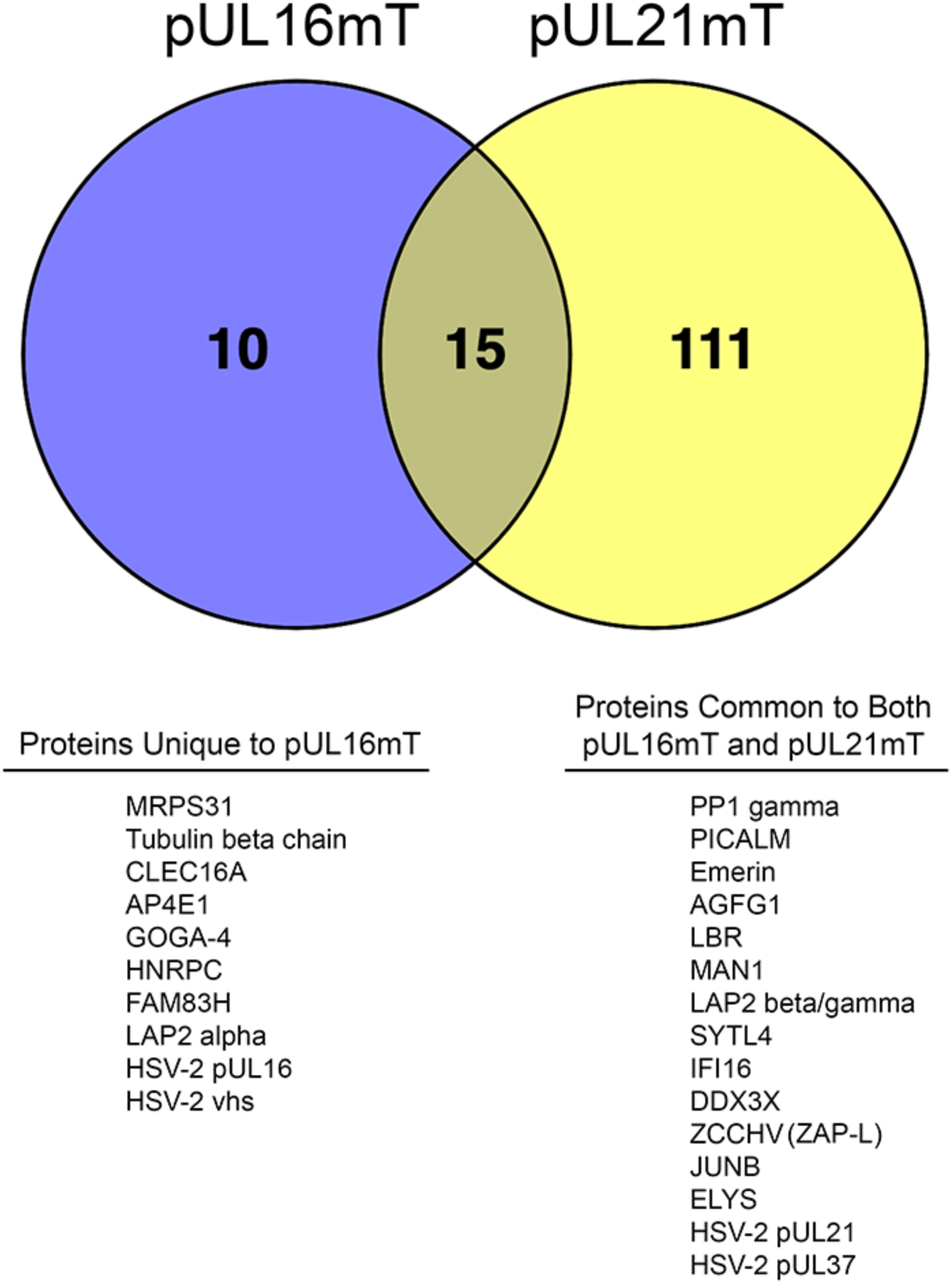
Comparisons of proteins in proximity to pUL16mT and pUL21mT during HSV-2 infection. **(A**) Venn diagram showing the numbers of proteins identified by BioID, meeting our stringent criteria, that were unique to pUL16mT (10), common to both pUL16mT and pUL21mT (15), and unique to pUL21mT (111). Lists of proteins unique to pUL16mT and common to both pUL16mT and pUL21mT are shown beneath the Venn diagram. Venn diagram was constructed using Venny 2.1 (https://bioinfogp.cnb.csic.es/tools/venny/).

In addition to their roles in the nuclear egress of nucleocapsids, pUL16 and pUL21 play important roles in the final envelopment of cytoplasmic nucleocapsids at membranes derived from the trans-Golgi network and/or late endosomes (22, 26, 46). Interestingly, phosphatidylinositol-binding clathrin assembly protein (PICALM), which plays a critical role in clathrin-mediated endocytosis and autophagosome formation (47), and synaptogamin-like protein 4 (SYTL4), which positively regulates exocytosis (48), were found in proximity to both pUL16 and pUL21. These findings may reflect their subcellular location during the late stages of virion maturation.

The antiviral molecules gamma-interferon-inducible protein 16 (IFI16) and the large isoform of zinc finger CCCH-type antiviral protein 1 (ZAP-L) were in proximity to pUL16 and pUL21 (Table 1, Figure 4, (39)). ZAP-L restricts the replication of a wide variety of viruses, including human cytomegalovirus (49), by binding to CG-rich viral RNAs and targeting them for degradation (50). Despite bearing a CG-rich genome, it has been reported that HSV-1 can replicate normally in the presence of ZAP-L (51), suggesting that ZAP-L is either ineffective against HSV-1 or that HSV-1 antagonizes ZAP-L activity. Studies by Sun and colleagues showed that phosphorylation of serine residues on ZAP-L by glycogen synthase kinase 3-beta (GSK3β) is critical for ZAP-L activity (52). It may be that complexes containing pUL16, pUL21 and PP1 gamma promote ZAP-L dephosphorylation/inhibition, thereby disrupting its antiviral activities. IFI16 is a well-known interferon-stimulated gene product that senses incoming herpesvirus genomes and promotes the expression of antiviral and pro-inflammatory cytokines that restrict virus replication and spread. IFI16 induction of cytokine expression in response to incoming HSV-1 genomes requires IFI16-driven liquid-liquid phase separation that is mediated through the phosphorylation of the IFI16 intrinsically disordered region by GSK3β and the cyclin-dependent kinase, CDK-2 (53). Similar to the scenario described above for ZAP-L, pUL16/pUL21/PP1 gamma complexes may promote the dephosphorylation of IFI16 to inhibit IFI16-mediated enhancement of cytokine expression.

Both pUL16 and pUL21 were found to be proximal to the viral inner tegument protein, pUL37, as well as the virion host shutoff (vhs) protein and tegument component, pUL41 (Table 1, Figure 4, (39)). As all four proteins are components of the HSV-2 tegument, their proximity may simply reflect their convergence during the late stages of virion assembly. However, the identification of pUL37 may have additional functional significance. Desai and colleagues reported reduced levels of pUL37 associated with nucleocapsids isolated from HSV-1 UL16, UL21, and UL16/UL21 double-mutant infected cells, suggesting that a pUL16/pUL21 complex directly, or indirectly, influences pUL37 loading onto nucleocapsids (25). Deletion of the UL37 gene is lethal for HSV-1 and mutant viruses demonstrated delayed nuclear egress of nucleocapsids and accumulation of non-enveloped nucleocapsids in the cytoplasm of infected cells, indicative of a secondary envelopment defect (54); phenotypes reminiscent of UL16 and UL21 deletion mutants (22, 23, 26, 46). This raises a question as to whether the deficiencies in secondary envelopment observed for UL16 and UL21 mutants are due to a failure of membrane-localized pUL16 to recruit pUL21-bound cytoplasmic nucleocapsids for secondary envelopment or due to deficiencies in the levels of pUL37 present on cytoplasmic nucleocapsids. Further definition of the roles of pUL16, pUL21 and pUL37 in the secondary envelopment of nucleocapsids is required to answer this question. Furthermore, pUL37 also functions in the early stages of HSV-1 infection to suppress the activity of an assimilated, nucleocapsid-associated kinesin microtubule motor. This enhances the efficiency of dynein-mediated nucleocapsid transport to the centrosome, where kinesin activity is then activated to deliver nucleocapsids to nuclear pore complexes for uncoating (13, 55). Thus, UL37 mutants demonstrate delayed uncoating due to a failure of nucleocapsids to efficiently reach the centrosome following viral entry. Interestingly, UL21 mutants derived from both HSV-1 and HSV-2 demonstrate delayed initiation of immediate-early gene expression after entry (37, 56). If reduced levels of nucleocapsid-associated pUL37 are a conserved feature between HSV species, the delayed gene expression phenotype may result from a failure of incoming nucleocapsids to be delivered efficiently to nuclear pore complexes. Understanding how pUL21 and pUL16 influence pUL37 recruitment to nucleocapsids is an important problem with implications for both early and late events in the HSV replication cycle.

Several proteins previously reported to interact with pUL16 were not identified as pUL16-proximal interactors in this study. These include the virion proteins pUL11, VP22 (UL49) and glycoprotein E (Us8) (26, 28, 30). In BioID experiments, a “bait” protein of interest (e.g. pUL16) is fused to a promiscuous biotin ligase, such as the third-generation enzyme mT, which is much more efficient and smaller than the BirA* and BioID2 enzymes (34). When cells expressing the mT-fusion are incubated with biotin, mT produces a short-lived reactive biotin species that becomes covalently linked to proximal lysine residues within an approximately 10 nm radius (35). Because biotin labelling depends on the presence of proximal lysine residues, proteins with low lysine content will be underrepresented, or absent, in BioID analyses. This explains the absence of pUL11, which contains no lysine residues, from our experiments. Peptides derived from both HSV-2 glycoprotein E and HSV-2 VP22 were identified in this study (Supplemental Data 1), but both proteins failed to surpass our stringent filtering criteria.

Ten of the 25 proteins identified in proximity to pUL16 during HSV-2 infection were not previously identified as being proximal to pUL21. These included the mitochondrial ribosomal protein S31 (MRPS31), and CLEC16A that regulates mitophagy (57). HSV-1 pUL16 is recruited to the mitochondrial surface, where it interacts with mitochondrial-associated membranes and localizes within mitochondria (32). In support of these findings, Li and colleagues identified the inner mitochondrial membrane protein adenine nucleotide transporter, ANT2, as an HSV-1 pUL16-interacting protein and determined that ectopic expression of pUL16 enhanced oxidative phosphorylation (33). Notably, ANT2 was identified as proximal to pUL16 in the present study but did not meet our filtering criteria (Supplemental Data 1). Taken together, these findings suggest that pUL16 is involved in the manipulation of mitochondrial function and turnover; however, additional studies examining the impact of pUL16 on mitochondrial physiology are warranted.

In summary, we have identified cellular and viral proteins in proximity to HSV-2 pUL16 in infected human keratinocytes. Leveraging our recent analysis of pUL21mT (39), together with this work, has given us insight into viral and cellular proteins proximal to both molecules, as well as unique interactors that may function in pUL21-independent activities. Intriguingly, these findings provide a nascent clue to the existence of different pUL21/PP1 complexes, some containing pUL16 and others not, that are PP1 isoform specific. Additional experimentation to confirm, or refute, the existence of different pUL21/PP1 complexes and, should they exist, to define their substrate specificities, is an exciting avenue of investigation for advancing our understanding of these fascinating HSV proteins.

## Materials and Methods

### Cells and viruses

Human keratinocytes (HaCaT), 293T, and Vero cells were acquired from the American Type Culture Collection (ATCC; Manassas, USA). HaCaT16 and Vero16 cells that stably express HSV-2 pUL16 have been described previously (21). HaCaT cells were maintained in Dulbecco’s modified Eagle’s medium (DMEM), supplemented with 10% fetal bovine serum (FBS; Wisent Bioproducts, Saint-Jean-Baptiste, Canada), 1% penicillin-streptomycin (Invitrogen, Burlington, Canada), and 1% GlutaMAX (Invitrogen), and grown at 37°C in a 5% CO_2_ environment.

The full-length infectious HSV-2 186 bacterial artificial chromosome (BAC), pYEbac373, was kindly provided by Dr. Yasushi Kawaguchi (University of Tokyo) and introduced into *Escherichia coli* (*E. coli*) strain GS1783 as described previously (Le Sage). Construction of the HSV-2 21-mT strain by *en passant* mutagenesis in pYEbac373 has been described previously (39). To construct 16-mT in pYEbac373, a PCR product was amplified using the pEP-Kan-S2 3x HAminiTurbo-in plasmid as a template using the forward primer 5’-TCGCGCGGGCCGTGGCCAGAGCCTCA TCGTCCGATTACAAAGCTAGCATCCCGCTGCTGAACGC-3’ and reverse primer 5’-CAGCCCCCC CCCCCGCCATGGCGGGGGGAAGCCTTACTGTTCAGTCGGCCCTGCTGAATTCC-3’. To facilitate recombination into the UL16 locus of pYEbac373, the forward primer contained 50 bp of sequence immediately upstream of the UL16 stop codon and in-frame with the mT open reading frame and the reverse primer contained a 50 bp sequence immediately downstream of the UL16 stop codon. The purified PCR product was integrated into pYEbac373 in strain GS1783 by *en passant* mutagenesis as described previously (37). To construct EGFP-mT, DNA sequences containing the HCMV major immediate promoter/enhancer driving the expression of an EGFP-mT fusion protein were cloned between 300 bp of HSV-2 (186) sequence found upstream of the HSV-2 Us2 open reading frame and 300 bp of HSV-2 sequence found downstream of the HSV-2 (186) Us2 open reading frame by Gibson assembly using the NEBuilder system, (New England BioLabs, Whitby, Canada). The resulting plasmid was linearized and co-transfected with purified HSV-2 (186) DNA into 293T cells. Infectious virus resulting from the co-transfection was used to infect monolayers of Vero cells and plaques were allowed to form for 72 hours. Plaques were screened for EGFP fluorescence using a Nikon TE200 inverted epifluorescence microscope. Fluorescent plaques were picked and purified three times, and isolates were tested for their ability to biotinylate infected cell proteins by fluorescence microscopy using Alexa Fluor 568-conjugated streptavidin (Invitrogen) at a dilution of 1:3,000.

### Antisera

Rabbit polyclonal antiserum against HSV pUL16 (29), a kind gift from Dr. John Wills (Pennsylvania State University), was used for western blotting at a dilution of 1:3,000. Rat polyclonal antiserum against HSV pUL21 (37) was used for western blotting at a dilution of 1:600. Mouse monoclonal antibody against BirA* (Novus Biologicals, Centennial, USA) was used for western blotting at a dilution of 1:500. Mouse monoclonal antibody against HSV-1 ICP27 (Virusys, Taneytown, USA) was used for western blotting at a dilution of 1:1,000. Mouse monoclonal antibody against β-actin (Sigma, St. Louis, MO) was used for western blotting at a dilution of 1:2,000. Horseradish peroxidase (HRP)-conjugated goat anti-rabbit IgG, HRP-conjugated goat anti-mouse IgG, and HRP-conjugated rabbit anti-rat IgG (Sigma) were used for western blotting at dilutions of 1:5,000, 1:15,000, and 1:80,000, respectively. Chicken polyclonal antiserum against HSV-2 pUL34 (20) was used for indirect immunofluorescence microscopy at a dilution of 1:500. Rat polyclonal antiserum against pUs3 (58) was used for indirect immunofluorescence microscopy at a dilution of 1:1,000. Alexa Fluor 568-conjugated streptavidin (streptavidin-568; Invitrogen) was used for indirect immunofluorescence microscopy at a dilution of 1:3,000. Alexa Fluor 647-conjugated donkey anti-rat IgG and Alexa Fluor 633-conjugated goat anti-chicken IgY (Molecular Probes, Eugene, OR) were used for indirect immunofluorescence microscopy at a dilution of 1:500.

### Preparation of whole cell extracts for western blot analysis

To prepare whole cell extracts for western blot analysis, 4×10^6^ HaCaT cells were infected at a multiplicity of infection (MOI) of 1, washed with cold phosphate-buffered saline (PBS), and scraped into 50-100 µL of PBS containing protease and phosphatase inhibitors (Roche, Laval, Canada), and transferred to microcentrifuge tubes containing 50 µL of 3X SDS-PAGE loading buffer. Transfected samples were treated with 250 units of benzonase nuclease (Santa Cruz Biotechnology, Dallas, USA) for 20 minutes at room temperature prior to the addition of 3X SDS-PAGE loading buffer. Cell lysates were passed through a 28 ½-gauge needle to reduce their viscosity, boiled for 5 minutes, and briefly centrifuged. For western blot analysis, 10-15 µL of cell lysates were electrophoresed through SDS-PAGE gels and separated proteins were transferred to polyvinylidene difluoride (PVDF) membranes (Millipore, Billerica, USA) and probed with appropriate dilutions of primary and secondary antibodies. Membranes were treated with Pierce ECL western blotting substrate (Thermo Scientific, Rockford, USA) and imaged using an Azure c300 Gel Imaging System.

### Virus replication analysis

4×10^6^ HaCaT cells were infected in triplicate at a multiplicity of infection (MOI) of 1 for replication analysis. After a one-hour inoculation period, cells were treated with low pH citrate buffer (40 mM Na citrate, 10 mM KCl, 0.8% NaCl) for 3 minutes at 37°C to inactivate extracellular virions, washed once with medium, and incubated at 37°C in complete medium, supplemented with biotin-depleted FBS with or without 50 µM of exogenous biotin. Medium and cells were harvested at 0, 6, 12, 24, and 48 hpi, subjected to three freeze/thaw cycles, briefly sonicated in a chilled cup-horn sonicator, and centrifuged at 3,000 rpm at 4°C to remove cellular debris. Titers were determined by plaque assay on Vero16 cells.

### Analysis of cell-to-cell spread of infection

Monolayers of HaCaT cells on 35 mm glass-bottom dishes (MatTek, Ashland, USA) were infected at various 10-fold dilutions required to yield well-isolated plaques and overlaid with complete medium containing 1% carboxymethylcellulose with or without biotin-depleted FBS. At 48 hpi, cells were washed with PBS three times then fixed and permeabilized with ice-cold 100% methanol, and stained for the HSV viral kinase pUs3 as described previously (58). Images of plaques (n=55 for each virus strain) were taken using a Nikon TE200 inverted epifluorescence microscope using a 10X objective and a cooled CCD camera. Plaque size was quantified by measuring plaque area using Image-Pro 6.3 software (Media Cybernetics, Bethesda, MD). Single infected cells were not included in this analysis.

### Indirect immunofluorescence microscopy

Infected cells were fixed at the indicated times post-infection with 4% paraformaldehyde in PBS for 15 minutes at room temperature. Cells were washed three times with PBS and permeabilized using 1% Triton X-100 in PBS for 10-30 minutes at room temperature. Following permeabilization, cells were washed three times with PBS/1% bovine serum albumin (BSA; PBS/1% BSA), incubated in primary antibody diluted in PBS/1%BSA for one hour at room temperature, washed three times with PBS/1% BSA, and incubated with appropriate Alexa Fluor conjugated secondary antibodies diluted in PBS/1% BSA for 0.5-1 hour at room temperature. After secondary antibody incubation, cells were washed three times with PBS/1% BSA, incubated with 0.5 µg/mL of Hoechst 33342 (Sigma) in PBS for 10 minutes at room temperature, and washed three times with PBS/1% BSA and stored at 4°C in PBS/1% BSA. Samples were imaged through a 60X (1.42 NA) oil immersion objective using an Olympus FV1000 confocal laser scanning microscope and FV10 ASW 4.01 software.

### Affinity purification of proteins biotinylated by pUL16mT

2.4×10^7^ HaCaT cells were infected with 16-mT at an MOI of 1. Following a one-hour inoculation period, cells were incubated at 37°C in complete medium, supplemented with biotin-depleted FBS and treated with 50 µM exogenous biotin for two hours at 4, 8 or 18 hpi. Cells were then washed with ice-cold PBS, harvested in 1 mL ice-cold PBS, and centrifuged at 500 x *g* at 4°C for 5 minutes. Biotinylated proteins were immobilized on streptavidin-Sepharose beads (GE Healthcare, Chicago, IL) according to the manufacturer’s instructions, with modifications based on a previously published protocol (59). Following incubation, bead-bound proteins were pelleted by centrifugation (Bound; B), and post-pull-down supernatants (S) were collected. A portion of each sample was retained prior to the addition of streptavidin-Sepharose beads (Input; I). Biotinylated proteins were visualized by western blot using SA-HRP.

### Sample preparation for BioID mass spectrometry

6×10^7^ HaCaT cells were infected with 16-mT or EGFP-mT in triplicate at an MOI of 3. Following a one-hour inoculation period, cells were incubated at 37°C in complete medium, supplemented with biotin-depleted FBS and treated with 50 µM exogenous biotin for two hours at 8 hpi. Cells were then washed with ice-cold PBS, harvested in 1 mL ice-cold PBS, and centrifuged at 500 x *g* at 4°C for 5 minutes. Biotinylated proteins were immobilized on streptavidin-Sepharose beads (GE Healthcare, Chicago, IL) according to the manufacturer’s instructions, with modifications based on a previously published protocol (59). Bead-bound proteins were digested with trypsin, and orbitrap/iontrap tandem mass spectrometry was performed by the Southern Alberta Mass Spectrometry Facility at the University of Calgary. Tandem mass spectra data were searched against a human protein database containing WT HSV proteins.

### Mass spectrometry data analysis

Tandem mass spectra data were searched against a human protein database containing WT HSV proteins using Mascot version 2.7.0 (Matrix Science, London, UK), assuming trypsin as the digestion enzyme. Mascot was searched with a fragment ion mass tolerance of 0.020 Da and a parent ion tolerance of 10.0 PPM. Scaffold version 5.3.3 (Proteome Software Inc., Portland, OR) was used to validate peptide and protein identifications. Peptide identifications were accepted if they could be established at greater than 95.0% probability by the Peptide Prophet algorithm (60) with Scaffold delta-mass correction. Protein identifications were accepted if they could be established at greater than 95.0% probability and contained at least one identified peptide. Protein probabilities were assigned by the Protein Prophet algorithm (61). Proteins that contained similar peptides and could not be differentiated based on MS/MS analysis alone were grouped to satisfy the principles of parsimony.

Previously published common BioID contaminants (42, 62) were removed from each dataset and proteins identified that met the following criteria were further analyzed: 1) a ≥ 95% protein and peptide threshold, 2) a minimum number of ≥ 3 identified unique peptides 3) ≥ 5% normalized percent coverage. Total spectral counts were normalized to the average spectral counts of endogenously biotinylated mitochondrial proteins (mitochondrial average): pyruvate carboxylase, mitochondrial, methylcrotonoyl-CoA carboxylase subunit alpha, and propionyl-CoA carboxylase alpha chain. For both minus-biotin and plus-biotin samples, normalization factors for each replicate were determined by dividing the highest mitochondrial average by each replicate’s mitochondrial average, and these factors were applied to all spectral counts within each replicate. Normalized spectral counts were determined by averaging values across three biological replicates, and minus-biotin averages were subtracted from the plus-biotin averages to generate the final normalized spectral counts. The average of the percent coverage of three biological replicates of the minus-biotin control samples was subtracted from the average of the percent coverage of the plus-biotin experimental samples to determine the normalized percent coverage.

### Statistical analysis

Graphical illustrations and statistical analyses were performed using GraphPad Prism version 10.4.2.

## Acknowledgements

This work was supported by the Canadian Institutes of Health Research (CIHR) Project Grants 162162 and 186187, Natural Sciences and Engineering Research Council of Canada Discovery Grants RGPIN/04249 and RGPIN/06733 and Canada Foundation for Innovation (CFI) award 16389 to BWB. SMH was supported in part by an Ontario Graduate Scholarship and Robert Sutherland and R. Samuel McLaughlin Fellowships from Queen’s University. The funders had no role in study design, data collection and interpretation, or the decision to submit the work for publication. We thank Dr. Laurent Brechenmacher from the Southern Alberta Mass Spectrometry Centre at the University of Calgary for expert assistance with mass spectrometry. We are grateful to the members of the Banfield laboratory for helpful comments on the manuscript.

## Notes

### Competing Interest Statement

The authors have declared no competing interest.

